# Specific degrader for fusion oncokinase kills tumors and is augmented by bimodal degrader-siRNA

**DOI:** 10.1101/2025.04.24.650501

**Authors:** Mahsa Shirani, Ronaldo Shaquille Bowie, Michael Tomasini, Ruth Hook, Denise Ng, Bassem Shebl, Barbara Lyons, Philip Coffino, Sanford M. Simon

**Affiliations:** Laboratory of Cellular Biophysics, The Rockefeller University; NY, NY 10065; Department of Chemistry and Biochemistry, New Mexico State University, Las Cruces, NM 88003

## Abstract

Oncogenic fusions of two native proteins are challenging to block without disturbing the parental proteins. A mRNA delivered peptide degrader, selectively eliminates the oncogenic fusions DNAJB1::PRKACA and ATP1B1::PRKACA in fibrolamellar carcinoma (FLC), cholangiocarcinoma and intraductal pancreatic neoplasm, without effect on native PRKACA. Specific degradation reflected structural properties of the fusion proteins. The degrader was lethal to FLC tumor cells in a preclinical model, with no effect on non-FLC cells. To enhance efficacy and speed, we modified the mRNA to simultaneously encode the degrader and an siRNA against the fusion transcript, reducing new protein synthesis which accelerated degradation of DNAJB1::PRKACA, offering a wide therapeutic index for mutation-driven disease.

## Main Text

Many cancers, such as FLC, Ewing’s sarcoma, chronic myelogenous leukemia and rhabdomyosarcoma, are driven by fusion oncoproteins ^2-4^. Since each fusion partner comes from a native protein that often has an essential function, it is important but difficult to target fusion oncoproteins while sparing native components. FLC is a usually lethal primary liver tumor of adolescents and young adults driven by an oncogenic fusion of the first exon of DNAJB1, a heat shock protein, to the bulk of the coding region of PRKACA, the catalytic subunit of protein kinase A (PKA) ^5-8^, forming DNAJB1::PRKACA. The fusion protein not only triggers the tumor, but elimination of the mRNA encoding DNAJB1::PRKACA with an shRNA ^9^ or an siRNA ^10^ is sufficient to stop growth and kill the tumor. The structure of the active sites of the oncogenic fusions DNAJB1::PRKACA (or an alternative fusion in cholangiocarcinoma and intraductal pancreatic neoplasm, ATP1B1::PRKACA) and that of native kinase are indistinguishable ^1^, making it difficult to find inhibitors specific to the fusions. Degraders such as molecular glues and proteolytic targeting chimera (PROTAC), have shown success in targeting proteins to the proteasome, usually using an E3 ligase to ubiquitinate the targeted protein. These have the potential for selective target degradation. However, it has proven problematic to selectively target the degrader to the fusion proteins and not the native components. We hypothesized that the distinguishing structural features of the native and fusion oncokinases might surmount the problem of identical structures in the kinase pocket, enabling selective degradation, even with ligands that bind equally to both proteins. We chose to design a degrader using a peptide from protein kinase inhibitor (PKI) ^11^, expressed as a fusion to an E3 adaptor. This empowers varying the properties of the peptide as well as freedom to optimize choice of the E3 pathway to match the pathology of the cancer. In FLC, the DNAJB1::PRKACA accumulates in the nucleus ^7^, suggesting the use of nuclear E3.

PKI was expressed as a fusion protein to Speckle-type POZ protein (SPOP), an E3 ligase that degrades its targets in nuclei ^12^. PKI-SPOP degraded DNAJB1::PRKACA and spared the native kinase, even though the PKI-SPOP bound equally to both. The DNAJB1 fragment was responsible for the preferential degradation by providing seven accessible lysines for the ubiquitination, thus targeting to the proteasome, and a flexible extended amino terminus ^1^ making the DNAJB1::PRKACA a good substrate for the proteasome. Eliminating the nuclear export signal in PKI trapped the degrader in nuclei and away from the cytosolic PRKACA. The modified PKI-SPOP degrader showed no toxicity to normal liver cells and selectively eliminated FLC tumors in mice. A degrader encoded by an mRNA enables a bimodal attack on both oncogenic protein and transcript, whereby a single mRNA molecule encodes both a degrader and an siRNA against the oncotranscript fusion junction. The combination, hitting both targets, was faster and more effective at eliminating the oncoprotein, compared to each single modality. Furthermore, a bimodal approach reduces the opportunity for tumor resistance or potential side effects, thus increasing the therapeutic index. A combined attack on the protein and its transcript shows potential for therapeutic development, offering hope for long-term help for patients.

## Results

### PKI-SPOP specifically degrades DNAJB1::PRKACA

We designed a degrader by replacing the native substrate binding domain of SPOP E3 ligase with the amino terminal 52 amino acids of PKI (PKI_1-52_)(Fig. 1A). SPOP was chosen due to its nuclear localization, where much of the DNAJB1::PRKACA is localized in FLC cells ^7^. Since FLC cells are oncogenically addicted to DNAJB1::PRKACA, we expressed the fusion construct in Huh7.5 cells ^13^. This allowed assessing degradation of DNAJB1::PRKACA independent of effects on cell viability.. Upon conditional expression of PKI_1-52_SPOP, there was effective degradation of DNAJB1::PRKACA, without a substantial effect on wild type PRKACA (Fig. 1B). Degradation was apparent within 3 hours of adding Dox, with the level of protein reduced two-fold within 12 hours (Fig. C, Supplement Fig 1) which parallels the time of expression and accumulation of the degrader.

**Fig. 1.**
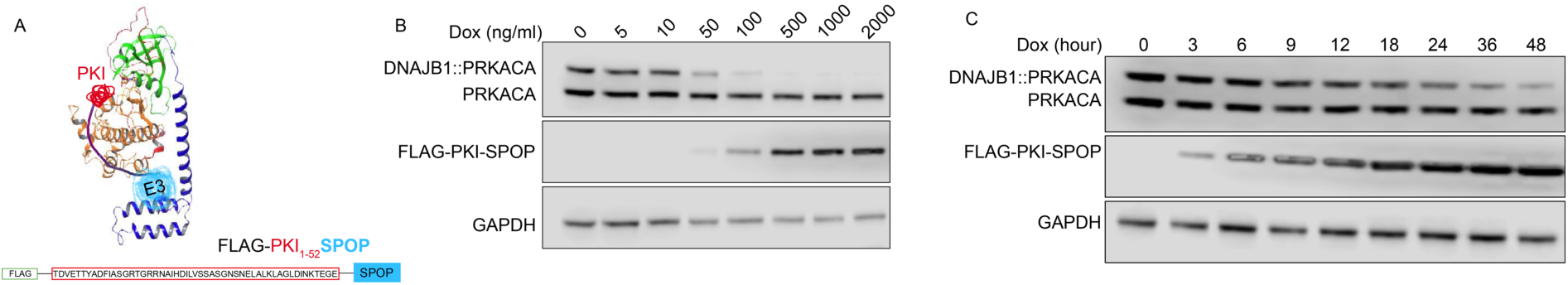
PKI-SPOP selectively degrades DNAJB1::PRKACA. **A) Schematic of PKI:SPOP.** DNAJB1::PRKACA fusion protein (modified from ^1^) with a cartoon of the PKI-SPOP. **B) Selective degradation of DNAJB1::PRKACA fusion.** Western blot analysis of Huh7.5 cells expressing endogenous PRKACA and transduced DNAJB1::PRKACA, treated with different concentrations of Dox for 72 h to induce expression of PKI_1-52_SPOP. **C) Time-dependence of degradation.** Western blot of DNAJB1::PRKACA in Huh7.5 cells after induction of expression of PKI_1-52_SPOP with 500ng/ml Dox, assessed by Western blot.

### PKI-SPOP mediated DNAJB1::PRKACA degradation requires ubiquitination and the proteasome

Degradation was dependent on proteasome proteolytic activity. The proteasome inhibitors MG132 (Fig. 2A) and bortezomib (Fig. 2B) rescued DNAJB1::PRKACA from degradation induced by expression of PKI_1-52_SPOP. Degradation of DNAJB1::PRKACA by PKI_1-52_SPOP was also rescued by impairing the ubiquitination cascade, using Tak-243, an inhibitor of E1 activating enzyme (Fig. 2C, D, E). Thus, degradation is ubiquitin dependent.

**Fig. 2.**
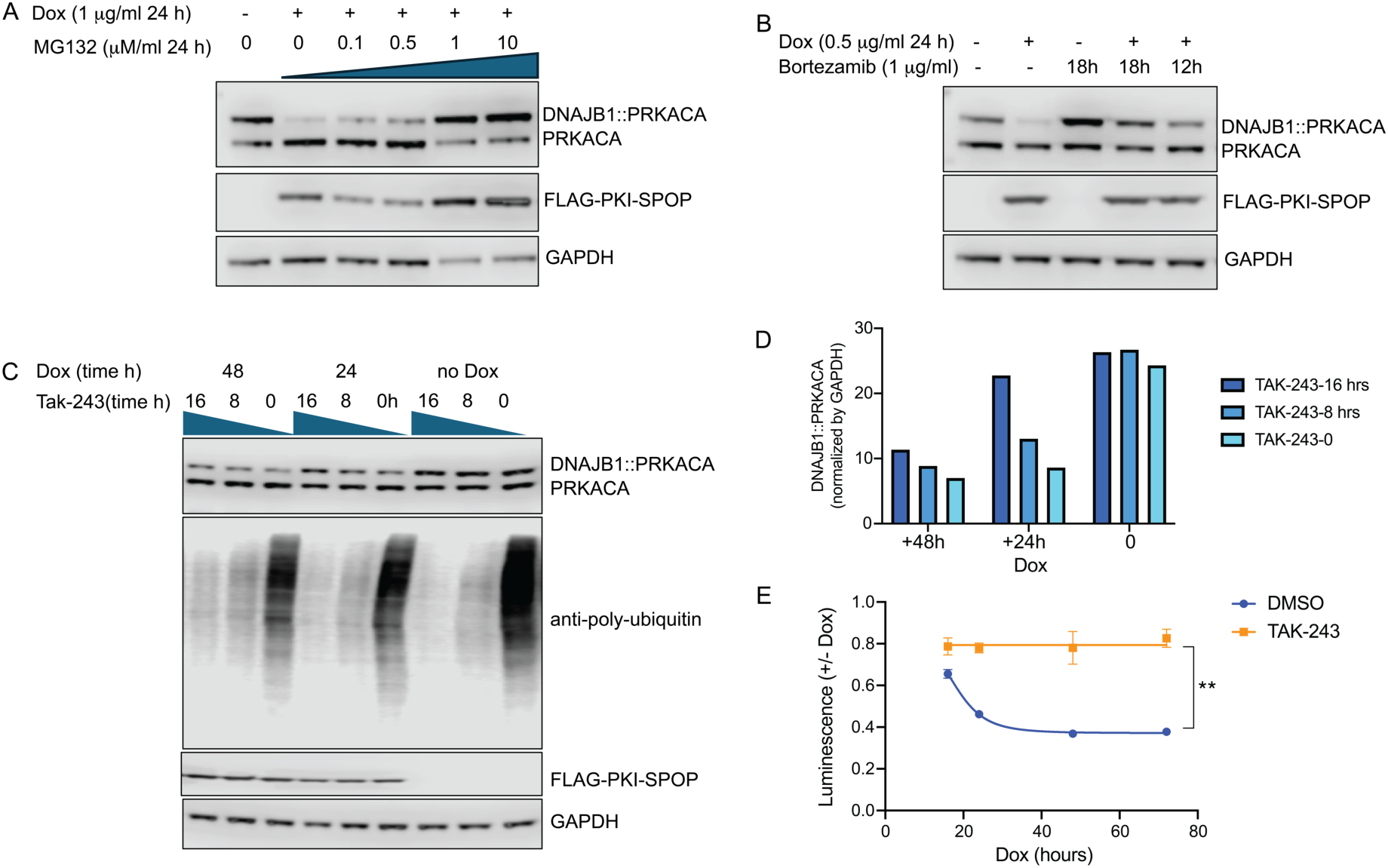
Degradation of DNAJB1::PRKACA upon induction of PKI_1-52_SPOP is dependent on the proteosome and ubiquitination. **A)** Levels of PRKACA and DNAJB1::PRKACA were monitored in Huh7.5 cells after induction of PKI_1-52_SPOP (24h) in the presence or absence of MG132. **B)** PRKACA and DNAJB1::PRKACA were monitored in Huh7.5 cells after induction of PKI_1-52_SPOP(24h). Cells were incubated with bortezomib (BZ) for 18 h or 12 h before harvesting. **C, D, E)** Dependence on E1 ubiquitination. To test the dependence of degradation on E1 activity, PKI_1-52_SPOP was induced in Huh7.5 cells for either 24 or 48h. Then the E1 inhibitor, Tak-243, was added for a duration of either 8hr or 16 hr prior to harvesting, or simultaneously with harvesting. **C)** Level of PRKACA and DNAJB1::PRKACA were monitored by Western blot. **D)** Quantification of the levels of DNAJB1::PRKACA, normalized to GAPDH from panel C. **E)** The level of Nano-luc-DNAJB1::PRKACA was monitored using luminescence in the absence and presence of Tak-243, at different time points subsequent to Dox induction and normalized to the no Dox counterpart.

### Dissection the mechanism(s) of selective degradation

Possible contributors to selective degradation include differential binding of the PKI-SPOP to DNAJB1::PRKACA and PRKACA, a specific topology between the SPOP and the DNAJB1::PRKACA, lysines accessible for ubiquitination in the DNAJB1 segment and flexibility of the DNAJB1 segment that would make the fusion a better substrate for the proteasome.

To assess whether PKI-SPOP differentially engages PRKACA and DNAJB1::PRKACA, we measured the kinase activity of each in the presence of increasing amounts of PKI-SPOP. The inhibition was indistinguishable, indicating similar affinity for both proteins (Fig. 3A).

**Fig. 3.**
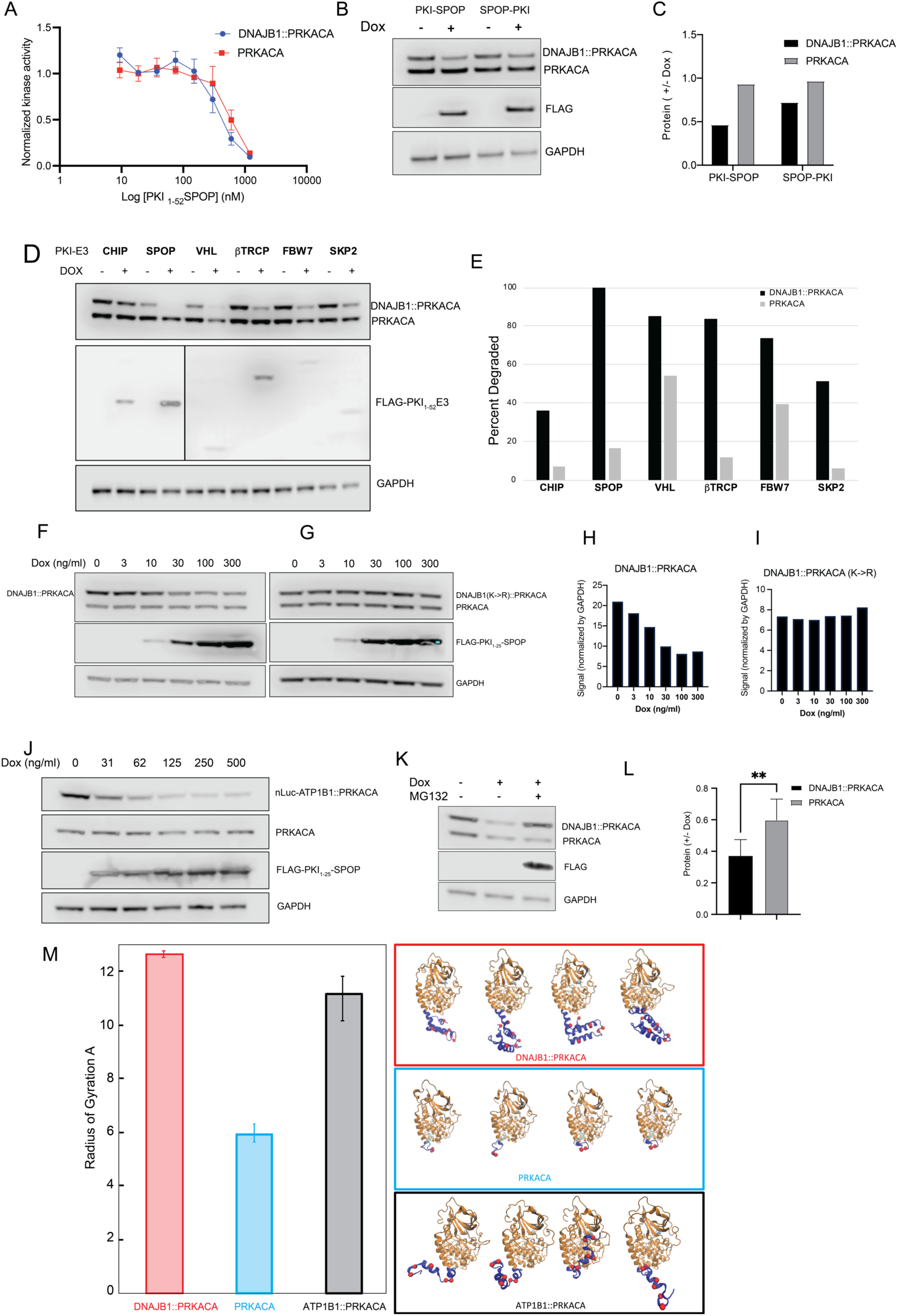
Specificity of PKI-SPOP for DNAJB1::PRKACA over PRKACA. A) Engagement of PKI-SPOP to target. PKI_1-52_SPOP similarly inhibits PRKACA and DNAJB1::PRKACA. Kinase activity was quantified at different concentrations of PKI_1-52_SPOP (n=3). **B, C) The effect of different PKI-E3 ligases on abundance of PRKACA and DNAJB1::PRKACA in Huh7.5 cells. B)** Western blot assay of the effects of expression of PKI-CHIP, PKI-SPOP, PKI-VHL, βTrCP-PKI, FBW7-PKI and SKP2-PKI. Each was analyzed in the absence (-) and presence (+) of 1μg/ml doxycycline (Dox), which induces the expression of the PKI-E3. (Quantification of DNAJB1::PRKACA and PRKACA bands in **B**, normalized by GAPDH are in Supplement 2). **C)** Fraction of DNAJB1::PRKACA or PRKACA remaining after adding dox. **D, E) Switching order of PKI and E3**. Huh7.5 cells were transduced with PKI_1-25_SPOP or SPOP-PKI_1-25_. **D)** The levels of DNAJB1::PRKACA and PRKACA were assayed by Western blot with an antibody that recognizes the shared carboxyl terminus. **E)** The ratio of each protein +/- dox plotted for each degrader. **F, G, H, I) Degradation of DNAJB1::PRKACA requires lysine in the amino terminus.** F) Western blot analysis showing degradation of DNAJB1::PRKACA by increasing concentration of Dox and F) abolition of degradation when all 7 lysines of its DNAJB1 domain are mutated to arginines (DNAJB1(KR)::PRKACA). **H)** Quantification of DNAJB1::PRKACA and **I)** DNAJB1(KR)::PRKACA (right), normalized by the level of GAPDH from western blots in A. **J) PKI_1-52_-SPOP degrades Nluc-ATP1B::PRKACA.** Increasing induction of degrader by Dox increased the selective degradation of Nluc-ATP1B::PRKACA with no effect on PRKACA as assessed by Western blot **K, L) Degradation upon direct targeting to proteasome**. PKI was expressed as a fusion to a peptide that binds to the PSMD2 subunit of the proteasome. Upon induction by Dox, there is a selective degradation of DNAJB1::PRKACA relative to PRKACA as seen by **K)** Western Blot and **L)** Quantification: the ratio of each protein +/- dox was plotted for each degrader. Quantification is from three independent experiments. **M) Dynamics of exon 1 of the fusion kinases and PRKACA.** Molecular dynamic simulations of DNAJB1::PRKACA (left) PRKACA (top), and ATP1B1::PRKACA (right) were used to calculate the radius of gyration of the amino terminus of each kinase (center bottom). Lysines in the amino terminal domains (blue) are marked by a red sphere. A space filling model of the MD simulations is in Supplement S3. The error bars are the standard deviation of three independent 1µs simulations for each kinase.

To test for the dependence of degradation on specific E3 complex configuration, we varied the linker length by shortening PKI or reversed the positions of binder and E3. DNAJB1::PRKACA was preferentially degraded relative to PRKACA with either PKI_1-25_SPOP or SPOP-PKI_1-25_ (Fig. 3B, C).

To further probe whether a specific E3 topology contributed to specificity, we replaced SPOP with different E3 ligases. PKI_1-52_ replaced the substrate binding domain of adapters of diverse E3 ligase complexes: von Hippel-Lindau (VHL), F-box/WD repeat-containing protein 7 (FBW7), S-phase kinase-associated protein 2 (SKP2), Speckle-type POZ protein (SPOP), beta-transducin repeat containing E3 ubiquitin protein ligase (βTrCP), and Carboxyl-terminus of Hsp70 Interacting Protein (CHIP) ^14,15^.

Most constructs (PKI-CHIP, PKI-SPOP, PKI-VHL, βTrCP-PKI, FBW7-PKI, SKP2-PKI) induced a decrease of DNAJB1::PRKACA (Fig. 3D,E, Supplement Fig 2). For some (SPOP, VHL), the levels of DNAJB1::PRKACA were lower before induction relative to others, which is likely the result of leakage of the Dox-inducible promoter. Some constructs led to preferential loss of DNAJB1::PRKACA (CHIP, SPOP, βTrCP, SKP2), while others (VHL, FBW7) resulted in greater loss of DNAJB1::PRKACA, but some loss of the wild-type PRKACA (Fig. 3D, E). These results suggest that retaining a precise spatial relation between kinase and E3 is not required for selectivity.

We next examined the role of lysines in degradation of the oncoproteins. The epsilon amines of lysines are the dominant targets for protein modification by the ubiquitination system. PRKACA has 2 lysines encoded by its first exon, and DNAJB1 has 7 lysines encoded by first exon. Changing the 7 lysines to arginines in the DNAJB1 domain of DNAJB1::PRKACA abolishes degradation of DNAJB1::PRKACA (Fig. 3F,G,H,I). Thus, selective degradation of DNAJB1::PRKACA was dependent on lysines in the DNAJB1 fusion.

If the specificity of degradation was a function of an amino terminal fusion that has lyines, then other fusions with lysines should also be susceptible. As PKI binds to the substrate binding site of the catalytic subunit, it should engage any fusion protein involving PRKACA. The fusion protein ATP1B1::PRKACA, found in some cholangiocarcinomas and in IOPN, has 6 lysines on the fusion adduct^16^. Induction of expression of PKI_1-52_SPOP led to degradation of nano-luciferase ATP1B1::PRKACA, as observed by western blot (Fig. 3J) or luminescence. (supplemental Figure 3). The results demonstrate that ATP1B1::PRKACA shares with DNAJB1::PRKACA the capacity to be targeted by a PKI-SPOP degrader.

To test whether E3-mediated ubiquitination of lysines was solely responsible for selective degradation of the fusion oncoprotein, we bypassed the ubiquitination cascade and directly targeted the kinases to the proteasome ^17^. We expressed PKI fused to a peptide that binds to PSMD2, a component of the 19S subunit that is near the opening of the proteasome ATPase pore, the site of substrate entry for degradation. While we observed high turnover of the degrader itself, we nonetheless observed 50% greater reduction of DNAJB1::PRKACA compared to PRKACA (Fig. 3K, L). This observation suggests that the DNAJB1::PRKACA fusion protein possesses intrinsic structural properties that make it more susceptible to proteasomal engagement and degradation, independent of an active ubiquitination pathway.

Degradation by the proteasome requires not only ubiquitination, but a substrate with a disordered or weakly ordered tail of sufficient size to initiate engagement by the proteasome ^18-20^. There is extensive structural information on PRKACA and DNAJB1::PRKACA ^1,21^, and we used AlphaFold ^22^ to predict the structure of the ATP1B1::PRKACA fusion (ribbon model in Fig. 3N, space-filling model in supplementary Fig. 4). With molecular dynamic simulations the amino terminal domain of native PRKACA has a much smaller radius of gyration than those of either DNAJB1::PRKACA or ATP1B1::PRKACA. This indicates that the amino terminus of both fusion oncoproteins is in a more extended conformation than that of PRKACA (Fig. 3N, lysines of each amino terminal domain marked in red). Even though the amino terminus of DNAJB1::PRKACA has more than twice as many amino acids as that of the ATP1B1::PRKACA, because it folds back on itself it has a similar radius of gyration. In general, the native PRKACA showed much less conformational flexibility than the two oncokinases in structural simulations. In analyzing the dynamics of the domain encoded by the first exon of each kinase, the two most occupied conformations of PRKACA accounted for over 90% of all the conformational space, whereas DNAJB1::PRKACA and ATP1B1::PRKACA showed more diverse conformational states (Supplemental Fig 4). Together, these results indicate that lysine residues in the DNAJB1 domain are essential for ubiquitination-driven degradation, while intrinsic structural properties of the fusion protein, such as extended amino-terminal flexibility, contribute to its greater susceptibility to proteasomal degradation.

### Variations of PKI that spare non-FLC cells

PKI_1-52_SPOP not only induces degradation DNAJB1::PRKACA, but also competitively inhibits the activity of both PRKACA and DNAJB1::PRKACA. Kinase inhibition is a potentially toxic effect independent of degradative targeting efficacy. As we expected, expressing PKI_1-52_SPOP in Huh7.5 cells, which harbor endogenous PRKACA but not the fusion protein, slowed cell growth (Fig. 4A).

**Fig. 4.**
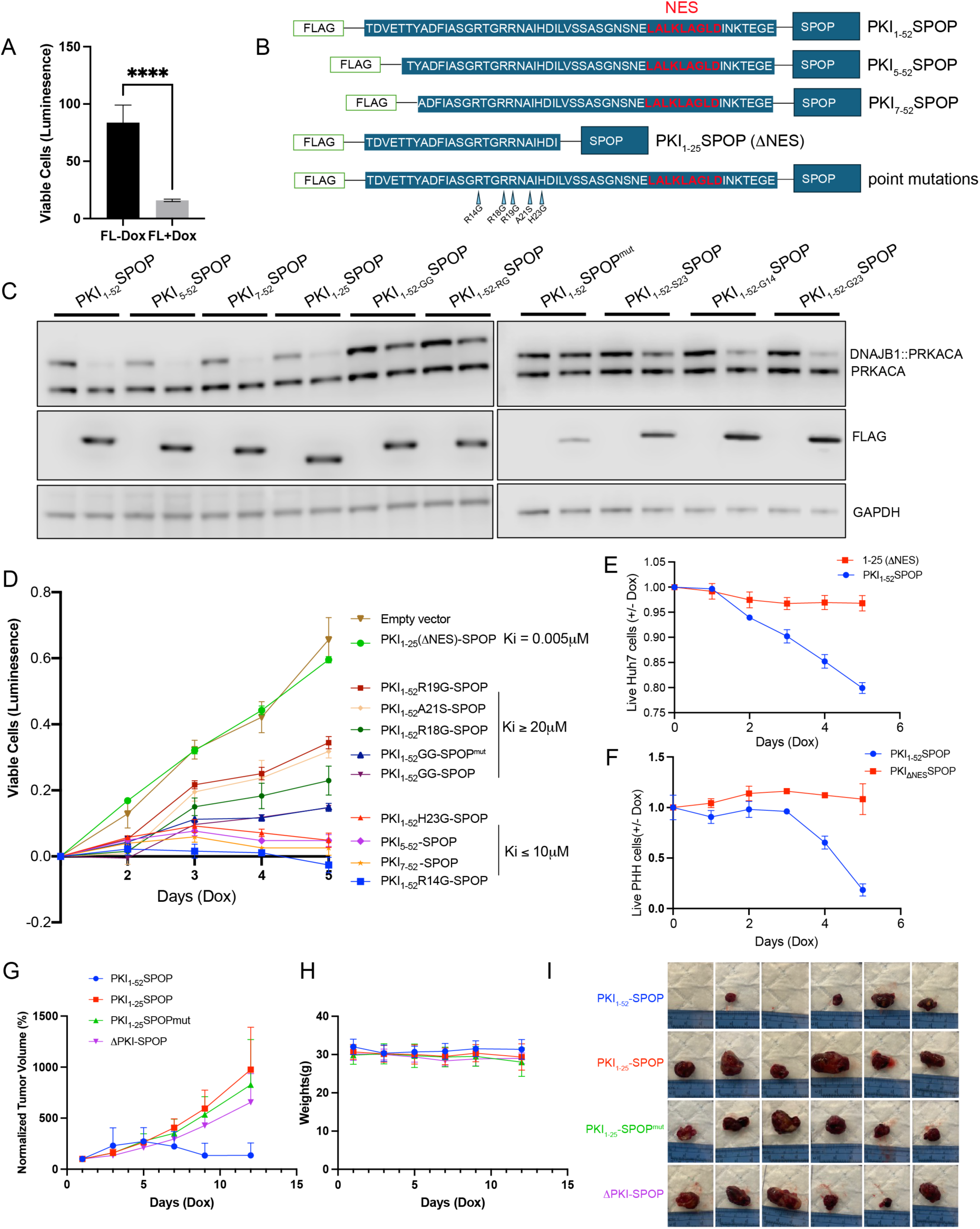
Effects of PKI-SPOP variants on degradation of DNAJB1::PRKACA and growth of HCC (Huh7.5) and FLC cells. **A)** Huh7.5 cells were induced to express PKI_1-52_SPOP for 5 days and live cells were quantified with cell titer glo in the absence and presence of Dox. **B)** Schematic representation of mutations to PKI. **C)** Western blot analysis of DNAJB1::PRKACA and PRKACA in the absence and presence of Dox in Huh7.5 cells transduced with different variants of PKI-SPOP. **D)** Growth of Huh7.5 cells that express endogenous PRKACA but not DNAJB1::PRKACA during 5 days of Dox treatment quantified with cell titer glo. **E)** Live cell count of Huh7.5 cells and **F)** PHH cells expressing PKI_1-52_SPOP and PKI_1-25_SPOP, normalized by cells not treated with Dox. **Expression of PKI-SPOP in Huh7.5 tumor G)** Tumor volume over time of Huh7.5 transduced with inducible PKI_1-52_SPOP, PKI_1-25_SPOP, PKI_1-25_SPOP^mut^ and ΔPKI-SPOP. Volumes were normalized to 100% on day 1. **H)** Weight of the mice over time. **I)** Tumors resected from mice in G.

To find modifications of PKI that could, when fused to SPOP, degrade DNAJB1::PRKACA but not slow the growth of Huh7.5 cells, we changed its sequence to reduce its binding affinity for the kinase catalytic subunit or to restrict its localization. (ΔNES) to nuclei (Fig. 4B). In normal cells, PRKACA is predominantly restricted to the cytosol, but in FLC cells the catalytic subunit is overwhelmingly in the nucleus ^7^. To reduce the affinity, we altered amino acids in PKI known to be involved in binding PRKACA. PKI missing its first 4 amino acids (PKI_5-52_SPOP) had a two-fold lower affinity for PRKACA and DNAJB1::PRKACA; PKI missing the first 6 amino acids (PKI_7-52_SPOP) has an ∼50 fold decrease in affinity; mutating the R14G (PKI_1-52-R14G_SPOP) causes a 1500 fold decrease and H23G (PKI_1-52-H23G_SPOP) a 500 fold decrease ^23^. These constructs remained able to degrade the fusion protein (Fig. 4C). Further decreasing PKI affinity by mutating either or both of two critical arginines, 18 and 19, to glycines decreases the affinity >5000-fold. These mutants produced less of an effect on the growth of Huh7.5 cells (Fig. 4D), yet maintained some degradation of DNAJB1::PRKACA (Fig. 4C). Changing alanine 21 to serine (which makes PKI a substrate rather than a pseudo-substrate and reduces its residence time on the catalytic subunit) lessened the inhibition of growth of Huh7.5 (Fig. 4D, PKI_1-52-A21S_), but also substantially decreased degradation of the fusion protein (Fig. 4C). Almost all variations of PKI tested decreased the growth rate in Huh7.5, thus risking an adverse effect on normal cells. Those variants of PKI with the highest binding affinity (≤10nM), inhibited growth the most. Those with lesser affinity (K_d_≥20 mM) did not slow growth as much.

In normal cells, PRKACA is predominantly restricted to the cytosol, but in FLC cells the catalytic subunit is overwhelmingly in the nucleus ^7^. To redirect the degrader to the nucleus we used a PKI that is truncated before its nuclear export signal (ΔNES), PKI_1-25_SPOP (Fig. 4B, NES highlighted in mauve). The PKI_1-25_SPOP had no effect on the growth of the Huh7.5 cells, providing a candidate for further development.

### PKI_1-25_SPOP in Huh7.5 and primary human hepatocytes

Huh7.5 and primary human hepatocytes (PHH) induced to express PKI_1-25_SPOP had the same growth rate as the empty vector control (Fig. 4D) and had the same growth rate as Huh7.5 (Fig. 4E) or PHH (Fig. 4F) in the presence and absence of Dox. Significantly, PKI_1-25_SPOP still effectively and selectively degraded DNAJB1::PRKACA over PRKACA (Fig. 4C). Therefore, PKI_1-25_SPOP was further characterized as a potential lead for degrader development.

We tested whether expression of these variants of PKI-SPOP, would have adverse effects *in vivo* on the growth of the HCC tumor Huh7.5, which does not express or depend on DNAJB1::PRKACA. Huh7.5 cells were stably transduced with plasmids encoding PKI_1-25_SPOP, PKI_1-25_SPOP^mut^ (lacking the three-box motif responsible for binding to Cullin E3); PKI_1-52_SPOP and SPOP without PKI (ΔPKI-SPOP). These were implanted subcutaneously in mice. When the tumor size was 100-300mm^3^, Dox was added to the diet to induce expression. The PKI_1-25_SPOP had no effect on growth of the Huh7.5 cells relative to expression of PKI_1-25_ with a mutant SPOP, or just SPOP alone (Fig. 4G, H, I). In contrast the PKI_1-52_SPOP, slowed the growth of Huh7.5 cells *in vivo* (Fig. 4G, H, I) consistent with its effects on the growth of Huh7.5 cells *in*

### PKI_1-25_SPOP in FLC

*vitro* (Fig. 4 D,E,F). Thus, a degrader based on PKI_1-25_ had no detectable deleterious effects on non-FLC hepatoma cells *in vivo*.

To assess the degraders in FLC tumor cells, we focused on PKI_1-25_SPOP, because it had no effect on viability of Huh7.5 cells but selectively and effectively degraded DNAJB1::PRKACA. FLC cells dissociated from patient derived xenograft (PDX) constitute a uniform population of FLC tumor cells, and the oncoprotein is their predominant form of the kinase. (In patient tumors the levels of the fusion and native kinase are more similar in bulktumor tissue because it is a mixed population of tumor, normal and stromal cells.) Cells were stably transduced to express the degrader FLAG-PKI_1-25_SPOP. We observed specific degradation of the fusion protein (Fig. 5A, middle lane), but the construct FLAG-PKI_1-25_SPOP^mut^ caused no degradation (Fig. 5A, right lane). We transiently transfected cells dissociated from PDX with the construct FLAG-PKI_1-52_SPOP. Using immunofluorescence, only cells expressing the degrader (yellow tag in Fig. 5B) showed a decrease of catalytic subunit (Fig. 5B).

**Fig. 5.**
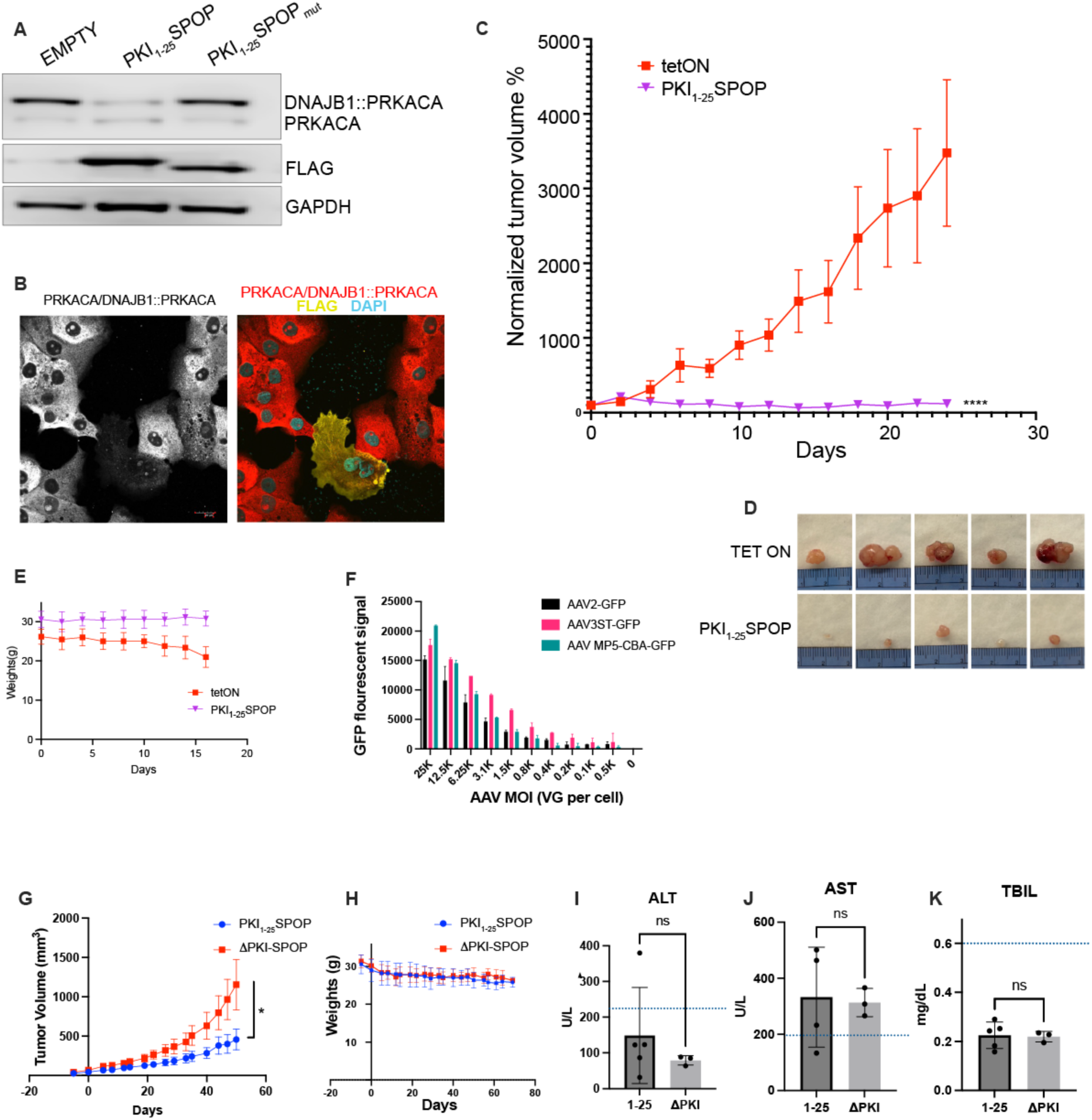
Expression of PKI-SPOP in FLC cells. **A) Western Blot analysis of FLC cells expressing PKI_1-25_SPOP or PKI_1-25_SPOP^mut^. B)Effect of PKI-SPOP on immunofluorescence of PRKACA and DNAJB1::PRKACA in FLC cells**. A subset of FLC-1 cells were transiently transfected with PKI-SPOP. An antibody against the carboxyl-terminus shows PRKACA and DNAJB1::PRKACA (red) and FLAG signal (from Flag-PKI-SPOP in yellow) and DAPI (cyan). **C) Effect of PKI-SPOP on growth of FLC tumors as PDX in mice**. Tumor volume over time for PDX of FLC-2 PDX transduced with inducible PKI_1-25_SPOP, and tet regulator. **D) PKI-SPOP reduces FLC tumor size.** Extracted PDX from mice in C. **E) Expression of PKI-SPOP has no adverse effects** on weight of the mice. **F) Efficacy of AAV in FLC cells.** Titration of efficacy of transduction by AAV strains AAV2, AAV3ST and AAV MP5 to FLC cells monitored by GFP fluorescence. **Delivery of AAV encoding PKI_1-25_SPOP or ΔPKI-SPOP (G, H, I, J, K). G) AAV with PKI_1-25_SPOP reduces tumor growth.** Effect of AAV delivery on growth of PDX of FLC-3, assessed by volume. **H) AAV delivery does not affect weight** of mice. **I-K) AAV delivery on mouse physiology.** ALT, AST and TBIL levels in the serum at the end of the study. Dotted line marks the upper limit of normal values in NSG mice.

### Degrading DNAJB1::PRKACA inhibits FLC growth *in vivo*

We tested whether expression of PKI-SPOP would affect the growth of FLC cells implanted as PDX in mice. FLC cells dissociated from PDX with inducible PKI_1-25_SPOP, were implanted subcutaneously in mice. When the tumor size was 10-40mm^3^, Dox was added to the diet to induce degrader expression. Tumor growth was completely abolished by the constructs in which SPOP was functional and PKI could bind the catalytic subunit (Figure 5C, D). PKI_1-25_SPOP, which lacks the nuclear export signal, was effective at blocking the growth of the FLC tumors (Fig 5C purple). The control group showed the greatest decrease in the weight of mice, consistent with a deleterious effect of unimpeded tumor growth on mouse health. In mice implanted with tumors and in which PKI_1-25_SPOP was expressed, there was no weight loss (Fig 5E). This indicates that PKI_1-25_SPOP was was protective of their health, through elimination of the tumor. Use of ΔPKI-SPOP did not affect tumor growth (Supplemental Figure 5A), and PKI_5-52_SPOP (slightly reduced affinity for PRKACA) showed inhibition of tumor growth, compared to PKI(_1-52_)SPOP^mutant^ (which blocks recruiting E1/E2) and PKI(GG)-SPOP with a reduced affinity for PRKACA (Supplemental Figure 5B).

To deliver the degrader by a viral vector we screened for adeno-associated virus (AAV) strains that could most efficiently confer GFP expression on FLC cells (Fig. 5F). We engineered AAV2 to express either PKI_1-25_SPOP or ΔPKI-SPOP (without PKI), driven by a CAG (cytomegalovirus early enhancer element, chicken beta-actin, rabbit beta-globin) promoter ^24^. FLC cells were implanted subcutaneously and AAV2 was delivered by tail vein injection. Tumor growth was slower with the AAV2 PKI_1-25_SPOP than with the negative control AAV2 SPOP (Fig 5G). There was no adverse effect of PKI on mouse weight (Fig. 5H) and no detectable liver toxicity, as measured by liver enzymes and metabolites, ALT, AST and TBIL, in the circulation (Fig. 5 I, J, K). Thus, a degrader based on PKI_1-25_ is lethal to FLC cells growing *in vivo*, with no detected effects on non-FLC hepatoma cells.

### siRNA against DNAJB1::PRKACA

Some potential and observed limitations on the efficacy of degraders include being overcome by high rates of new protein synthesis, mutations in the target protein and mutations in the protein degradation pathway. Each of these could limit the ability of the degrader to get sufficiently complete and rapid decrease of the oncoprotein to trigger cell death. We tested the efficacy of a attack on the protein and its encoding mRNA by combining the degrader PKI_1-25_SPOP with an siRNA against the transcript fusion junction. When DNAJB1::PRKACA was expressed from the DNAJB1 promoter (e.g., Fig. 1), there was clear efficacy of PKI_1-25_SPOP. However, its efficacy was difficult to observe by Western blot when the fusion protein was expressed from the stronger EF1α promoter (Fig. 6A). A combination degrader/antisense could potentially overcome tumor escape by overexpression of DNAJB1::PRKACA. A bimodal approach would potentially have other benefits. Since the antisense hits the fusion junction and the degrader hits the catalytic pocket, it would require two different forms of resistance to escape the combined therapy. Further, each will have different potential side effects. The combination would increase the therapeutic window.

**Fig. 6:**
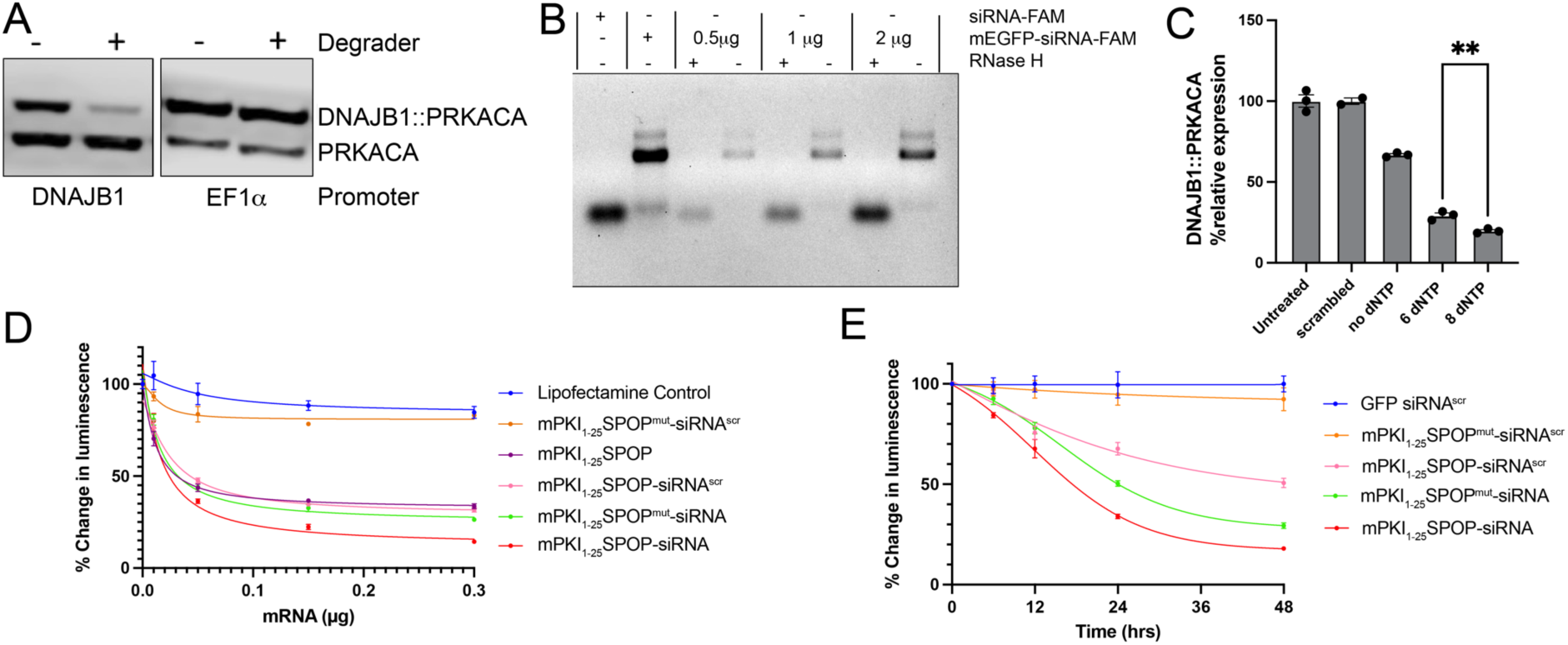
Multimodal degrader and antisense. **A) Efficacy of degrader is affected by the level of oncoprotein.** Left: PKI_1-25_SPOP was effective at eliminating most DNAJB1::PRKACA with no effect on PRKACA when the oncoprotein is driven by the DNAJB1 promoter. Right: However, when DNAJB1::PRKACA is driven by the stronger EF1a promoter, the same amount of PKI_1-25_SPOP is insufficient to eliminate most of the oncoprotein. The PRKACA in each sample is the native protein driven from the native promoter. The Western blot on the right was not exposed as long, to avoid saturating the DNAJB1::PRKACA. **B) DNA/RNA hybridized to the mRNA results in RNase H cleavage between the polyA tail and RNAi sequences.** The far left lane is just the siRNA. All the remaining lanes have the mRNA for mEGFP and the siRNA. The second lane from the left is the full length mEGFP-siRNA. The next three pairs of lanes have variable amounts of the mRNA for mEGFP-siRNA added with or without RNase H. **C) Effect of dNTP on RNase H processing.** The number of dNTP in the DNA that hybridizes to spacer between the polyA tail of the degrader and the siRNA was varied. Huh7.5 cells were transfected with the hybrid mRNA-siRNA and processing of the RNA was assessed. **D) Efficacy of degrader and antisense**. Huh7.5 cells were transduced to stably express DNAJB1::PRKACA-nanoluciferase from the DNAJB1 promoter. mRNA encoding the bimodal degrader/siRNA (PKI_1-25_SPOP-siRNA, red), PKI_1-25_SPOP^mut^-siRNA (which inactivated SPOP, green), PKI_1-25_SPOP-siRNA^scr^ (with the siRNA sequence scrambled, pink), or PKI_1-25_SPOP (purple), or and PKI_1-25_SPOP^mut^-siRNA^scr^ (orange) were transcribed in vitro. The Huh7.5 cells expressing DNAJB1::PRKACA-nanoluciferase were transfected with varying concentrations of mRNA. Luminescence was recorded 48 h after transfection. **E) Time course of degrader and antisense.** mRNA encoding the biomodal degrader/siRNA (PKI_1-25_SPOP-siRNA red), PKI_1-25_SPOP^mut^-siRNA (green), PKI_1-25_SPOP-siRNA^scr^ (pink), PKI_1-25_SPOP^mut^-siRNA^scr^ (orange) or GFP-siRNA^scr^ (blue) were transcribed in vitro and cells treated as in **D**.

We took advantage of the nucleic acid (mRNA) formulation of our degrader to design a single molecule that encoded both the degrader and an siRNA against the fusion junction of the oncotranscript ^10^. A combined single molecule would facilitate delivery, as well as future preclinical and clinical implementation. After the polyA tail of the degrader we added RNAi sequences at the 3′ end of mRNA. We adopted a strategy of “Combined hybrid structure of siRNA tailed IVT mRNA”, whereby a structure of DNA/RNA is hybridized to the mRNA between the polyA tail and RNAi sequences ^25^. This should result in the siRNA being cleaved from the mRNA by RNase H. We tested this approach using a GFP followed by an siRNA to the DNAJB1::PRKACA oncotranscript. mRNA was transcribed *in vitro*, incubated with the short RNA/DNA hybrid, and then transfected into Huh7.5 cells. The siRNA was cleaved from the mRNA in the cells (Fig. 6B). Increasing the number of dNTP in the DNA, thus increasing the length of the recognition site for RNase H, increased the efficacy of cleavage of the siRNA (Fig. 6C).

We next tested the effects PKI_1-25_SPOP combined with an siRNA targeting the fusion junction of DNAJB1::PRKACA in Huh7.5 cells (Fig. 6D). mRNA encoding the bimodal degrader/siRNA (PKI_1-25_SPOP-siRNA), or PKI_1-25_SPOP, PKI_1-25_SPOP-siRNA^scr^ (with the siRNA sequence scrambled), or PKI_1-25_SPOP^mut^-siRNA, and PKI_1-25_SPOP^mut^-siRNA^scr^ were transcribed *in vitro* and transfected into Huh7.5 cells expressing a DNAJB1::PRKACA-nanoluciferase. The PKI_1-25_SPOP (just the degrader), PKI_1-25_SPOP-siRNA^scr^ (the degrader with a scrambled siRNA) and the PKI_1-25_SPOP^mut^-siRNA (the siRNA with a nonfunctional degrader) all produced a similar decrease on the amount of DNAJB1::PRKACA. The combination of the degrader and siRNA was more efficacious at decreasing DNAJB1::PRKACA at all concentrations. Neither the vehicle control or the PKI_1-25_SPOP^mut^-siRNA^scr^ affected DNAJB1::PRKACA.

We examined the time course of DNAJB1::PRKACA-nanoluciferase degradation induced by the degrader, the siRNA or the combination of the two (Fig. 6E). The degrader alone (PKI_1-25_SPOP-siRNA^scr^ pink) was the slowest. Presumably, this is because the mRNA must first be translated into protein. The siRNA (PKI_1-25_SPOP^mut^-siRNA, green) was faster. The fastest, and most effective was the combination, PKI_1-25_SPOP-siRNA (red). The combination of the degrader and siRNA also most effectively reduced DNAJB1::PRKACA which was driven from the EIF1a promoter (Supplemental figure 6).

## Discussion

Many cancers are driven by fusion proteins. Several questions arise in considering whether these fusion oncoproteins are favorable therapeutic targets. Do they merely trigger the tumor or is their continued action required to drive the tumor? Second, are the tumors oncogenically addicted: do they die when the fusion oncoprotein is eliminated or instead revert to a normal phenotype? The latter would necessitate continuous treatment of the tumor and exacerbate the risk of mutational escape from the therapeutic. Our data imply favorable answers to these two questions for FLC. A third critical question is whether the fusion protein can be targeted without impairing either of the contributing parental proteins. Either or both native proteins may have important cellular roles. If so, one must find ways to selectively eliminate the fusion protein.

One approach for specificity is the use of antisense technology. The junction of the fusion transcript is unique. Therefore, specificity can be obtained by siRNA, shRNA or anti-sense oligonucleotides targeted to the junction to restrict continued synthesis of the oncoprotein. We have used shRNA to destroy DNAJB1::PRKACA mRNA and have shown that FLC is oncogenically addicted to the fusion oncoprotein ^9^. The use of siRNA can also eliminate the fusion protein ^10^. This powerful ensemble of transcript-directed techniques must overcome problems of *in vivo* delivery of antisense nucleic acids to the tumor cells and into their cytosol. Additionally, a method that impairs synthesis may be limited by long-lived oncogenic proteins, which persist after their production is stymied. The DNAJB1::PRKACA protein lasts for days after elimination of its mRNA ^9^. With a slow reduction of protein levels, there is a risk that cells start to normalize rather than die of oncogenic addiction.

A second approach for specificity is the use of degraders. Degraders must overcome the problem of restricting proteolytic degradation to the fusion protein, and not to its parental partners. If achievable, this is a powerful approach. Degraders must also overcome continued synthesis of new protein from mRNA. Here we demonstrate we can achieve specificity of degradation of the fusion oncoprotein, with no effect on the parental partners and overcome the problem of new synthesis by combining, in the same molecule, a degrader with an siRNA.

We demonstrate that selective degradation does not require selective binding. Indeed, we developed an agent that binds similarly to DNAJB1::PRKACA and PRKACA, but nonetheless selectively degrades the fusion (Fig. 1B). We show this is because its amino-terminal extension, by being flexible and lysine rich, enhances ubiquitination and proteasome engagement. Relieving the requirement for selective binding could enable identification of degraders for other proteins. For a targeting agent we used a peptide fragment from PKI that binds selectively and with high affinity to the catalytic subunit of PKA at its substrate binding site ^11^. The peptide was expressed as a fusion to various adapters of different E3 ligases, most of which yielded preferential degradation of DNAJB1::PRKACA over PRKACA. A larger PKI fragment of 52 amino acids was lethal to FLC tumors. While the PKI_1-52_SPOP did not degrade native PRKACA, it inhibited the activity of PRKACA, rendering it deleterious to non-FLC cells. While PRKACA is usually in the cytosol, we recently observed that the overexpressed catalytic subunits in FLC cells are predominantly in the nucleus ^7^. Thus, nuclear colocalization of target and degrader could facilitate selective effects on FLC over other cells. In this work, we found one of the most effective degraders was based on SPOP, which has a nuclear import signal and acts in the nucleus ^26^. To trap PKI-SPOP in the nucleus, we removed the nuclear export signal from PKI. This provided an effective and selective degrader of DNAJB1::PRKACA over PRKACA. Most importantly, degradation of the fusion protein with PKI_1-25_SPOP eliminated FLC cells *in vitro* and FLC tumors in mice, while showing no adverse effects in non-FLC cells.

Degradation induced by PKI-SPOP requires lysines in the adduct contributed by the DNAJB1 moiety of the fusion protein, depends on ubiquitination, and requires proteasome degradative activity. It has been observed that the activity, and specificity, of some degraders is very sensitive to the precise configuration of the lysines on the target protein, the degrader, the particular E2 ubiquitin conjugating enzyme and E3 ligase ^27-29^. A specific spatial configuration of the ternary complex that delivers ubiquitin and extends ubiquitin chain conjugates might account for the specificity of degrading the fusion over the native kinase, as seen with some degraders of Bcr::Abl over the Abl kinase ^30^ or with p38 MAP kinase orthologs ^31^. However, we show here instead that the observed degradation specificity is unlikely to depend on the specific configuration of the degradation complex and E3. We observe preferential degradation of DNAJB1::PRKACA over PRKACA with multiple E3 ligases (Fig. 3, D,E). Additionally, we observe specific degradation with a variety of lengths of linkers to the E3 ligases, ranging from 25 amino acids (PKI_1-25_) to 52 amino acids (PKI_1-52_). Further, SPOP-based degraders were efficacious whether the E3 was positioned at the amino or the carboxyl end of the targeting peptide (Fig. 3B, C). These findings suggest that the specificity we observe does not depend on a specific configuration of the ubiquitination complex, but is instead a robust property of the substrate itself.

There are two plausible possibilities. First, the extension and flexibility of the oncoprotein fusion domain may facilitate its engagement by the ATPase ring of the proteasome. Proteasome substrate degradation is enhanced by an extension of adequate size and flexibility for initiation of proteasome entry ^18,19,32^. The bulk of the native PRKACA catalytic subunit is tightly folded. In contrast, the N-terminal adducts of DNAJB1::PRKACA and ATP1B1::PRKACA extend from the PRKACA core and are more dynamic and less compact than the corresponding element of native PRKACA (Fig. 10) (Tomasini et al., 2018). The greater size and more dynamic conformations of the oncogenic adducts likely promote their engagement by the proteasome ^33^ and may contribute to their differential degradation. A second possibility is that the lysines within the fusion domain may be favorably positioned for ubiquitination and proteasome association. Within the ternary complex, orientation of the E3 relative to potential target lysines has been shown to be critical for some selective degraders ^29,31^. The number and distribution of lysines along the fusion domain may affect the efficacy and the position of the ubiquitin chain adducts, both of which matter for degradation ^34^. The domain of ATP1B1 that is fused has 6 lysines and that of DNAJB1 has 7 lysines. In contrast, there are only two lysines in the corresponding N-terminus of PRKACA. Further, the amino terminus of PRKACA is myristoylated and anchored close to the kinase core (Fig. 4) ^1,35^. We show that that direct targeting to the 19S subunit of proteasome, bypassing the E3 ligase and thus the requirement for ubiquitination, still yields preferential degradation of the fusion protein (Fig 3L, M), although not as selective as when targeting is through the E3 pathway. Thus, the amino terminal extension contributes some preferential degradation, which is further enhanced by the ubiquitination of lysines in the amino terminus. The present findings broaden the possibilities for designing degraders for other fusion proteins that share the favorable structural characteristics of the FLC fusion kinases.

The degrader used here is encoded by an mRNA. Small molecule degraders are limited in design by the variety of E3 ligases that can be targeted, usually restricted to VHL and CRBN. An advantage of the mRNA encoded approach is the ability to test a wide variety of E3 ligases (Fig. 3D, E). This allows finding E3 ligases that are most efficacious in a particular tumor line. Using mRNA to broaden the choice of E3 ligases also facilitates a design that reduces the likelihood of mutational escape. By selecting an E3 that the tumor relies on, the risk of escape through loss of that E3 pathway is reduced. Additionally, concomitant use of multiple degraders using different E3s further reduces the potential of mutational escape. A further advantage of an mRNA-encoded degrader that incorporates a peptide binder: the peptide can be modified to fine tune degrader localization or target binding affinity, thereby promoting specificity, and avoiding off-target toxicity. Our degrader started from a naturally occurring peptide that binds with high affinity and high specificity. This can be generalized to other diseases with existing peptide-display libraries. Additionally, the rapid advance of protein language models facilitates the design of peptides that selectively target fusion or mutated oncoproteins^36-38^. Another major advantage of an mRNA-encoded degrader is the ability to generate an siRNA on the same molecule. The siRNA reduces the ability of the cell to synthesize new oncoprotein, thereby increasing both the efficiency (Fig 6B) and speed (Fig 6C) of degradation of the oncoprotein. The combination of the two into a single molecule reduces the need for separate regulatory testing of safety and efficacy of the degrader, the antisense, and the combination. This could be critical for rare cancers where the number of patients available for a clinical trial is often a limitation.

The combination of hitting both the mRNA and the protein offers the further advantage of reducing the chance of escape by diverse mechanisms specific to the protein, its transcript or to the E3 degradation apparatus. A tumor could escape the antisense with a mutation to the codons encoding the fusion junction or with a mutation that affects the binding of the PKI to the catalytic site, or the protein degradation pathway ^39-42^. The combination reduces or delays mutational escape. Additionally, either the antisense, or the degrader, could have low levels of off target effects. The combination of the two, each with different potential off target effects, increases the specificity and improves the therapeutic window.

Efficiently targeting an mRNA-encoded degrader to tumor cells can be challenging. Fortunately, recent years have seen an explosion of technologies for delivering mRNA to different organs and different cells using lipid nanoparticles ^43-45^. This approach must be tuned for each tumor and is one that offers potential for FLC. FLC cells are similar to normal hepatocytes and retain expression of the asialoglycoprotein receptor, enabling targeting by the sugar GalNac on the surface of lipid nanoparticles, an approach we have shown to work with siRNA ^10^. FLC cells also have relatively normal expression of the LDL receptor and other proteins that have been successfully used for targeting liver cells. The similarity of FLC to normal liver may instead be a liability, for the liver can act as a sink, diverting most of the lipid nanoparticles from tumor. Moving forward, tumor-specific targeting might be facilitated by identifying proteins uniquely expressed on the surface of tumor cells. This approach may be particularly important for eliminating micrometastases.

## Acknowledgments

We would like to thank the fibrolamellar patients, and their caregivers, through their contributions to The Fibrolamellar Registry, to our Fibrolamellar Tissue Repository, through work at the bench and contributions in too many ways to enumerate.

## Funding

NIH/NCI P50CA210964 (SMS)

NIH/NCI U54CA243126 (SMS, MS, PC)

NIH/NCI R01CA248507 (SMS)

NIH/NCATS UL1 TR001866 from a Clinical and Translational Science Award (CTSA) (SMS)

KOODAC is funded by Cancer Research UK (CGCATF-2023/100019), Institut National Du Cancer (INCa), and KiKa (Children Cancer Free Foundation) through Cancer Grand Challenges (SMS, MS, MT, PC)

Supported in part by grant, Center for Basic and Translational Research on Disorders of the Digestive System through the generosity of the Leona M. and Harry B. Helmsley Charitable Trust (SMS);

The Rally Foundation 23CN18 (SMS)

## Author Contributions

Conceptualization: MUS, RSB, RS, PC, SMS

Methodology: MUS, RSB, MT, DN, BS, SMS

Investigation: MUS, RSB, MT, RH, DN,

Visualization: MUS, RSB, MT, SMS

Funding acquisition: MUS, PC, SMS

Project administration: SMS

Supervision: BAL, PC SMS

Writing – original draft: MUS, SMS

Writing – review & editing: MUS, SMS, MT, BS, PC, SMS

## Data and Materials availability

All data are available in the main text or the supplementary materials

## Supplementary Materials

### Methods

#### Plasmids

To generate Tet-On 3G bidirectional inducible plasmids, gene fragments for FLAG-PKI_1-52_SPOP_167-374_, FLAG-PKI_1-52_VHL_152-213_, FLAG-PKI_1-52_CHIP_128-303_, FLAG-CRBN _2-320_PKI_1-52_, FLAG-βTrCP_2-263_ PKI_1-52_, FLAG-FBW7_2-293_PKI_1-52_ and FLAG-SKP2_2-147_PKI_1-52_ were synthesized by Azenta and cloned into pRetroX-TRE3G-mCherry-CMV promoter (TaKaRa Bio 631188 Cat #641188) using NEBuilder HiFi Assembly (New England Biolabs Cat # E2621S). Q5 site directed mutagenesis (NEBiolabs Cat # E0554S) was used for further mutations to truncate PKI or SPOP (constructs PKI_5-52_SPOP, PKI_7-52_SPOP, PKI_1-25_SPOP, PKI_7-25_SPOP, PKI-SPOP^mut^ (lack the three box motif responsible for binding to Cullin) (ΔAAEILILADLHSADQLKTQAVDFIN) or to replace E3 with KLHDC2 targeting peptide (SPPPMAGG) or PSMD2 targeting peptide MC2 (RDKPLHRYVGFQC). QuickChange Lightning (Agilent Technologies Cat #210518) was used to generate point mutations on PKI (PKI_R19G_SPOP, PKI_A21S_SPOP, PKI_R18G_SPOP, PKI_R18G_R19GSPOP, PKI_R14G_SPOP, PKI_R23G_SPOP). To generate nano-luc-DNAJB1::PRKACA, nanoluc was synthesized by Azenta and cloned on the 5’ end of DNAJB1::PRKACA (pcrDNA3.1 Addgene plasmid #100891). Nano-luc-ATP1B1::PRKACA was generated (Azenta) by replacing the DNAJB1::PRKACA on the nano-luc DNABJ1::PRKACA. The DNAJB1::PRKACA with the lysines changed to arginine on the DNAJB1 adduct were synthesized by Azenta.

Note on numbering conventions: In many articles, the amino acids of PKI do not count the initiating methionine, so threonine, the second amino in the genomic sequences, is counted as 1. To facilitate comparison with the rest of the literature we assume this convention and did not count the initiating methionine in the numbering.

#### Cell culture

Human hepatocellular carcinoma Huh-7.5 cells (Blight et al., 2002), human embryonic kidney HEK293T cells (ATCC Cat#CRL-3216) and immortalized PHH, Hu1545 ^46^, gift of Rory Smoot, were grown in Dulbecco’s Modified Eagle’s Medium supplemented with L-glutamine, sodium pyruvate (DMEM, Gibco, Cat #2904457), and 0.1 mM non-essential amino acids (Gibco Cat #11140-050) and 10% fetal bovine serum (FBS, Sigma, Catalog #F4135) in humidified incubators at 37°C and in a 5% CO_2_ and 5% O_2_atmosphere. The additional cell lines used in this work were generated by transducing these cells with retroviral and lentiviral vectors (see below) with the plasmids described above.

#### Plasmid transfection

Plasmid transfection of FLC cells was done using FuGENE® HD (Promega, Cat #E2311) according to the manufacturer’s instructions.

#### Retrovirus and Lentivirus particle production

Low-passage HEK293T cells at 60-80% confluence were co-transfected using either Lipofectamine^TM^ 2000 (Invitrogen, Cat #11668019) or polyethyleneimine (PEI MAX, Kyfora Bio, Cat#24765) and a 5:5:1 ratio of a retroviral or 25:5:1 ratio of lentiviral plasmid, a suitable GagPol plasmid, and a VSV-G plasmid, respectively. 11 μg of total DNA were used to transfect a 10-cm dish, in a total of 10 mL of media. For higher titer, 88 μg of total DNA were used to transfect a 15-cm dish in a total volume of 20 mL of media. Virus-producing HEK293Ts were maintained in DMEM+ supplemented with 10% FBS and 1% non-essential amino acids. Virus-containing media was collected at 24 h, (and at 48 h and 72 h, as needed) post transfection, filtered through a 0.45 μm filter (EMD Millipore, Cat #SLHPR33RS), and used to transduce target cells immediately, after short storage at 4°C, or frozen at -80°C. For virus concentration, pooled virus-containing media was mixed at a ratio of 3:1 with LentiX Concentrator (Takara Cat # 631232), cooled at 4°C, then centrifuged for 45 min at 4°C and 1,500 x g. The viral pellet was resuspended into OptiMEM (Thermo Fisher Sci, Cat #31985070) to achieve a 100-fold concentration and stored at -80°C. Infections of Huh-7.5-derived cell lines were done in DMEM+ supplemented with 10% FBS, and 10 μg/mL Polybrene (EMD Millipore, Cat#CR-1003-G), for 6 to 16 h. Infections of FLC cells were done in RPMI (Gibco Cat#11875-119) supplemented with 10%FBS. Infected cells were selected using related antibiotics. Selection was started two days after transduction. Antibiotic selection of transduced Huh-7.5-derived cell lines, PHH and FLC cells was done with 3 μg/mL puromycin (Invitrogen, Cat # A1113803), 600 μg/mL G418 (Thermo Scientific, Cat # 10131035), or 6 μg/mL blasticidin(Sigma, Cat # 15205).

#### Cell viability assay

Huh7.5 cell lines were plated in triplicate with a density of 5k-10k per well in black 96-wells plates in DMEM supplemented with 10%FBS, NEAA and 500 ng/ml of doxycycline. Growth of Huh7.5 cells and PHH were measure either daily using PrestoBlue HS (Thermo Scientific, Cat # A13261) or as a dead end assay using CellTiter-Glo reagent (Promega, Cat # G7572), according to the manufacturer’s instructions.

#### Treatment of cells with compounds

The following chemicals were added to cell media at concentrations and times stated in the results: MG132 (Sigma, Cat # M7449), bortezomib (Selleck Chemicals, Cat # S1013), TAK-243 (MLN7243, Selleck Chemicals, Cat # S8341).

#### Luminescent assay

Cells expressing the Nano-luc tag on the amino terminus of ATP1B1::PRKACA and on the carboxyl terminus of DNAJB1::PRKACA, were plated down in white 96 wells plate. NanoLuc® Luciferase (Promega, Cat #N1110) was used to measure the luminescent signal according to the manufacturer protocol.

#### Mice

All procedures were carried out under approved animal use protocols from The Rockefeller University (New York, NY) Institutional Animal Care and Use Committee approval (#20027-H). NOD-*scid* IL2Rgamma^null^, NOD-*scid* IL2Rg^null^, NOD *scid* gamma (NSG) mice were obtained from Jackson Laboratories (Strain #:005557) and housed in the specific pathogen-free immune core at The Rockefeller University animal facility. The mice were maintained on a 12-hour light/dark cycle, provided an amoxicillin-supplemented diet and had free access to food and water. Both male and female mice were used for patient-derived xenograft (PDX) passaging and *in vivo* studies. At the time of tumor implantation, mice were between 1 and 2 months old. Their health and tumor growth were monitored at least twice per week.

For the in vivo studies, mice were anesthetized using isoflurane on mice. 300K to 1millon cells were injected subcutaneously with 1:1 ratio of RPMI and Matrigel (Corning Cat#356231). The mice were fed with 1,000 ppm doxycycline food (inotive, Cat#TD.120658) when a detectable size of tumor formed. Tumor size was measure by caliper measure two or three times a week.

#### Tumor dissociation

Mice bearing tumors subcutaneously, were euthanized and tumors resected. Patient-derived xenografts (PDXs) were cut and placed in 50-mL Falcon tubes containing RPMI medium, 2% penicillin/streptomycin, collagenase D (Sigma Cat #11088882001), Protease (Sigma Cat #P5147) and DNase I (Sigma Cat #10104159001). Digestion was performed at 37°C with rotation until the tissue was fully dissociated (Benchmark Scientific Roto-Therm Cat# H2024). All subsequent steps were carried out on ice or at 4°C. The digested tissue was filtered through a 200-μm strainer (Pluriselect Cat#43-50200-03), using a syringe plunger to push through any remaining fragments, and then through a 100-μm strainer (Thermo Fisher Scientific, Cat #352360). The cells were centrifuged at 321 × g for 5 minutes at 4°C, followed by red blood cell ysis with a brief 10-second exposure to of water, after which phosphate buffered saline (PBS) (Gibco Ref #14190-144) was added. Cells then underwent mouse cell depletion according to the manufacturer’s protocol (Miltenyi Biotec, Cat#130-104-694). Cells were then counted and plated on plates that we coated with collagen (Sigma Cat #C3867-1VL) for transduction.

#### AAV gene transfer

Recombinant AAV expressing PKI_1-25_SPOP and SPOP under TBG promoter were made by Vector Biolab. AAV strains expressing GFP (AAV2-GFP, AAV3ST-GFP and AAV MP5-CBA-GFP) were gift from Ype De Jong. For *in vitro* work, FLC cells were dissociated as described previously and plated down in black 96 wells plates. AAVs were added to the cells in different titers with 2-fold serial dilutions (0.5K-25K VG per cell) in triplicates. GFP signal was measured after 48hours using the plate reader. For the *in vivo* studies, FLC cells injected subcutaneously into mice (n=5 per condition) and when a detectable size of tumor was observed, AAV vectors were injected through the tail veil twice with two days in between each injection and at 5 x 10^11^ vector genome per mouse.

#### Western blot analysis

Cells were lysed in ice-cold RIPA buffer (Alfa Aesar, J63306-AP) supplemented with cOmpleteTM EDTA-free protease inhibitor cocktail (Roche, Cat #11697498001), PhosSTOP (Roche, Cat #301210) and DNaseI (Sigma Cat #10104159001) for 20 min. Cell lysates were centrifuged at 20,000 g, 4°C for 10 min and supernatants were collected. Protein concentration was measured using the BCA protein assay kit (Pierce, Cat#23227). 10 to 50 μg of total protein was run on 4-12% Bis-Tris gels (Thermo-Fisher, Cat#164820), transferred onto nitrocellulose membranes using the iBlot (Invitrogen, Cat #AV23002), and blocked for 1-2 hours at room temperature in 5% milk in Tris-buffered saline supplemented with 0.1% Tween-20 (Fisher Cat #BP337) (TBST) buffer. Blots were probed with the primary antibodies against PRKACA (Cell Signaling Technology Cat# 5842, RRID:AB_10706172, 1:3000), GAPDH (GeneTex Cat# GTX627408, RRID:AB_11174761, 1:5000) or Ubiquitin (Sigma-Aldrich Cat# 05-944, RRID: AB_3668631, 1:2000), for overnight at 4°C in blocking buffer, followed by the secondary antibodies anti-mouse (Sigma-Aldrich Cat# A9917, RRID:AB_258476, 1:10,000) IgG and goat anti-rabbit IgG (Sigma-Aldrich Cat# A0545, RRID:AB_257896, 1:10,000-1:25,000) for 1 hour at room temperature. HRP tagged primary antibody against FLAG (Sigma-Aldrich Cat# A8592, RRID:AB_439702) was used to detect FLAG signal. Amersham ECL detection kit (GE Healthcare, Cat #115300) was used to develop the blot. Imaging was done using LI-COR Blot Scanner (C-DiGit). Blots were stripped with Restore Plus Western Blot Stripping buffer (Thermo, Cat #46430) to probe for multiple proteins.

#### Microscopy: immunofluorescence

FLC cells were plated on Cultrex Poly-D-Lysine (biotechne, Cat #3439-100-01) treated and collagen coated MatTek dishes at 37C in 5% pCO_2_. 48 hours following transfection, cells were fixed with 4% (wt/vol) paraformaldehyde (Electron Microscopic Sciences, Cat #15711) in PBS for 10 min (room temperature). They were washed 3 times for 5 min with PBS. Cells were blocked and permeabilized for 1 hr in blocking buffer (2.5% normal donkey serum (Jackson Lab Cat #017-000-121), 2.5% normal goat serum (Sigma, Cat #G9023), 0.1% vol/vol) Triton X-100 (Sigma Cat #T9284), and 1% BSA (Sigma Cat#A9418). Primary antibodies against PRKACA (Cell Signaling Technology Cat# 5842, RRID:AB_10706172, 1:200) and FLAG (Sigma-Aldrich Cat# F3165, RRID:AB_259529, 1:200), were then added in blocking buffer and incubated overnight at 4C in a humid chamber. Cells were then washed 3 times for 5 min in PBS and then incubated with secondary anti-mouse antibody (Thermo Fisher Scientific Cat# A48286, RRID:AB_2896351, 1:1000) and goat anti-Rabbit (Thermo Fisher Scientific Cat# A-11036, RRID:AB_10563566, 1:1000) in blocking buffer for 1–2 hr at room temperature. Dishes were dried and mounted with SlowFade Diamond Antifade Mountant (Invitrogen, Cat #P36961). Cells were imaged via confocal microscopy (Olympus FluoView FV3000, UPLXAPO 60× oil immersion objective, numerical aperture: 1.42).

#### Protein Purification

##### DNAJB1-PRKACA and PRKACA

Human DNAJB1::PRKACA and PRKACA were cloned into pET151 (InVitrogen Cat# K15101). Purification protocol was adapted from ^47^. Proteins were expressed in Escherichia coli (E. coli) BLD21 (DE3) pLysS. Protein expression was induced at 18°C overnight by 0.4 mM isopropyl β-D-thiogalactoside (IPTG) (Research Products International Cat #156000-25.0) at an OD_600_ of 0.8. Cells were resuspended in TMN50 (50 mM Tris pH 7.0 RT, 50 mM NaCl, 2 mM MgCl_2_) supplemented with cOmpleteTM EDTA-free protease inhibitor cocktail tablets (Roche), PhosSTOP (Roche) and NaATP (1 mM) (TCI Chemicals Cat #A0157-25G) and lysed using a high pressure homogenizer (Avestin EmulsiFlex-C3). Lysate was clarified at 20,000xg for 20 min. The clarified lysate was incubated with Affi-gel10 resin (Bio-RAD Cat #1536099) conjugated to the peptide PKI_5-24_ (synthesized by Thermo Fisher). Bound protein was washed with TMN250 buffer (50 mM Tris pH 7.0 RT, 250 mM NaCl, 2 mM MgCl_2_) supplemented with NaATP (1 mM) and eluted in elution buffer (50 mM Tris, pH 7.0 RT, 50 mM NaCl, 2mM EDTA and 200 mM L-arginine). Protein was concentrated and stored in storage buffer (TMN50 + 10% glycerol).

##### PKI-SPOP

FLAG tagged PKI-SPOP was expressed in primary human hepatocyte (PHH). Cells were lysed in 50mM Tris pH7.4, 150 mM NaCl, 1mM EDTA and 0.5% triton X-100. Cells lysate was clarified by centrifugation (14,000 x g for 10–15 min at 2–4°C). ANTI-FLAG® M1 Agarose Affinity Gel (Sigma Cat#A2220) was used to purify the FLAG-PKI_1-52_SPOP according to the manufacturer protocol.

##### Molecular Dynamics

MD simulations were performed for the fusion proteins DNAJB1::PRKACA and ATP1B1::PRKACA as well as the native PRKACA. Each protein was simulated with ATP, 2MG^2+^ ions and residues 5 to 24 of PKI (PKI5-24). The DNAJB1::PRKACA systems were built using chain A of the crystal structure, PBD ID: 4WB7^48^ in which the protein is bound to ATP, several Zn^2+^ ions, and PKI5-24. Zn^2+^ions associated with nucleotide were converted to Mg^2+^ ions, with other Zn^2+^ions removed. The PRKACA systems were built from the crystal structure PDB ID: 4DFX ^49^. This structure is a mouse PRKACA structure in complex with the ATP mimic AMP-PNP, 2 Mg^2+^ions and the 20 residue peptide SP20, a variant of PKI_5-24_ with two mutations (N20A and A21S) which convert PKI_5-24_ from inhibitor to substrate. The structure also includes the myristoylation at the N-terminus as well as an additional mutation K7C. Residues 7, 32, 34, 39, 44, 65, 124, 348 of PRKACA were mutated to convert the mouse PRKACA structure to human PRKACA. Additionally, residues 20 and 21 were mutated to convert SP20 to PKI. AMP-PNP was mutated to ATP and the 2 Mg^2+^ions were retained. The ATP1B1::PRKACA systems were built using Alphafold3 ^50^ to contain the ATP1B1::PRKACA fusion protein, PKI, ATP, and 2Mg^2+^ ions. All structures were phosphorylated at S139, T197 and S338 (PRKACA numbering). Structures were processed using the Protein Preparation Wizard in Maestro (Maestro 10 (2016) Schrödinger, Inc., Portland, OR), solvated in a rhombic dodecahedron water box with SPC waters, and 150mM salt (sodium and chloride ions). Simulations were performed using the Desmond MD Package ^51^ using the OPLS4 force field ^52^. Each system was subject to energy minimization using the steepest decent method. An initial 100 ps Brownian Dynamics simulation at constant volume and a temperature of 10 K with heavy atoms constrained was performed.

Subsequent equilibration included a 12 ps simulation at constant volume and at 10 K with heavy atoms restrained, followed by a 12 ps simulation at constant pressure with heavy atoms restrained, and finally a heating simulation in which the restraints were gradually relaxed and the system heated to 300 K over 24 ps. Production simulations were performed for 1 μs with system snapshots saved every 25 ps.

##### Radius of Gyration Calculations

Helices E (Residues 140–160) and F (Residues 217–233) (PRKACA numbering) where used to align trajectories of each individual simulation. The radius of gyration (Rg) of the was calculated using residues 1-14 for PRKACA, 1-31 for ATP1B1, and 1 to 69 for DNAJB1 using the VMD molecular graphics software package ^53^. The Rg for a given kinase was taken as the average of three independent 1 μs simulations.

##### Clustering of Conformations

For each kinase simulations were aggregated to generate distinct clusters of conformational states. Helices E (Residues 140–160) and F (Residues 217–233) (PRKACA numbering) where used to align trajectories to the initial crystal structure. Analysis was performed on the protein backbone atoms first using the GROMACS ^54^ analysis tool ‘gmx rms’ to compute the root-mean-square-deviation between all structures in the trajectory. The GROMACS utility ‘gmx cluster’ was then used to cluster the structures using a cutoff of 8 Å and the gromos clustering methodology ^55^. Representative structures were taken as the structure with the smallest average RMSD distance to every other member of the cluster.

**Supplement Figure 1:**
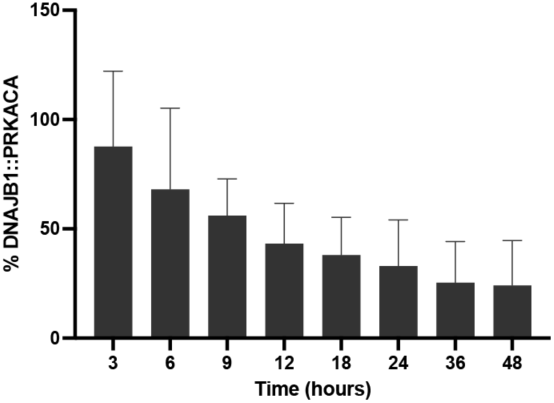
Quantification of DNAJB1::PRKACA from Western blot in Figure 1C. Error bars are standard deviation, n=2.

**Supplemental Figure 2.**
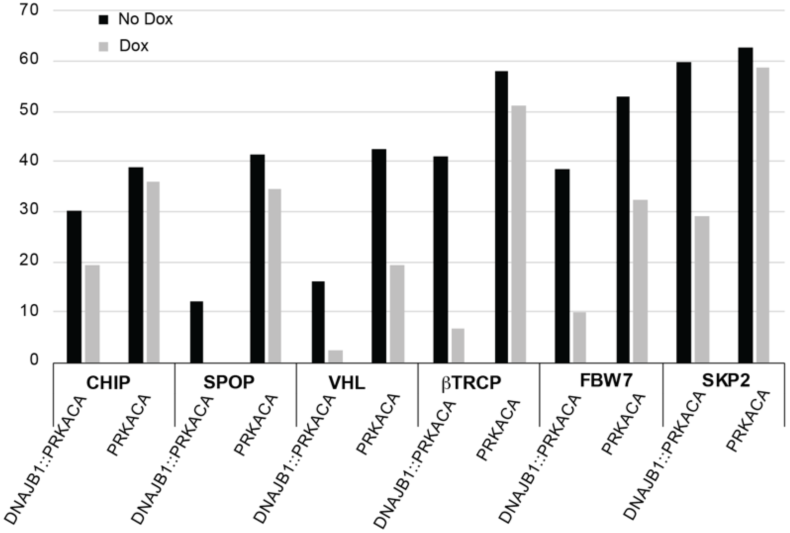
Quantification of DNAJB1::PRKACA and PRKACA bands from Figure 3D, normalized by GAPDH

**Supplement figure 3:**
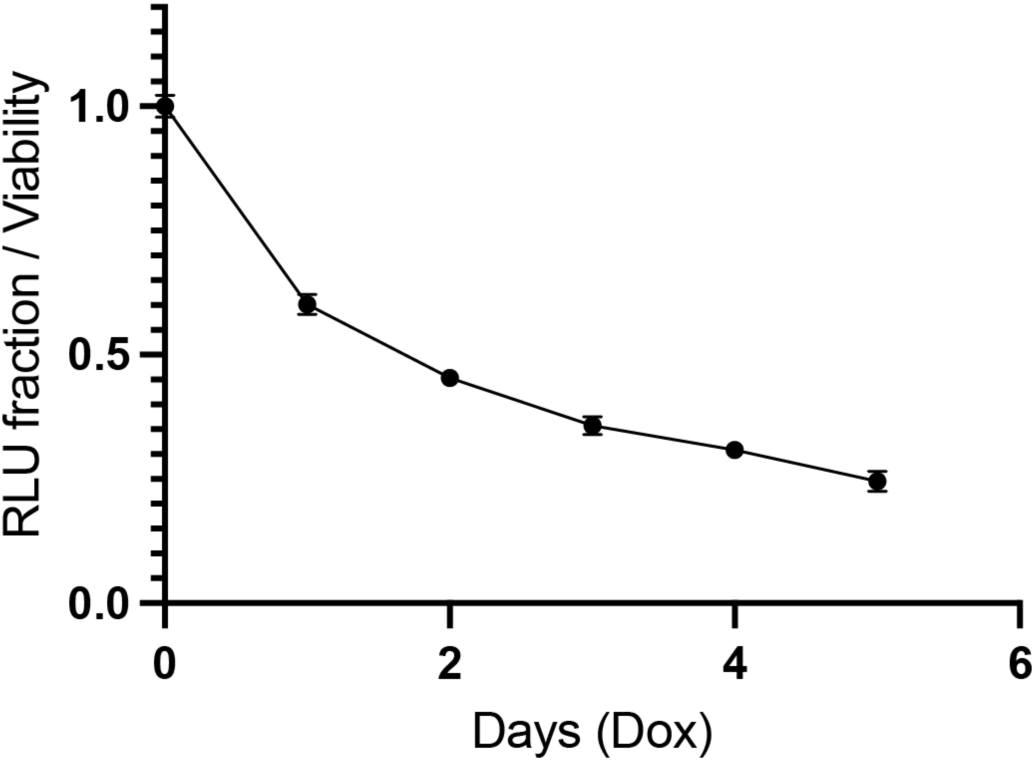
Degradation of nano-luciferase-ATP1B1::PRKACA at time points after Dox treatment, measured by luminescence and normalized for cell viability, as measure by Cell-titre Glo^TM^.

**Supplemental Figure S4.**
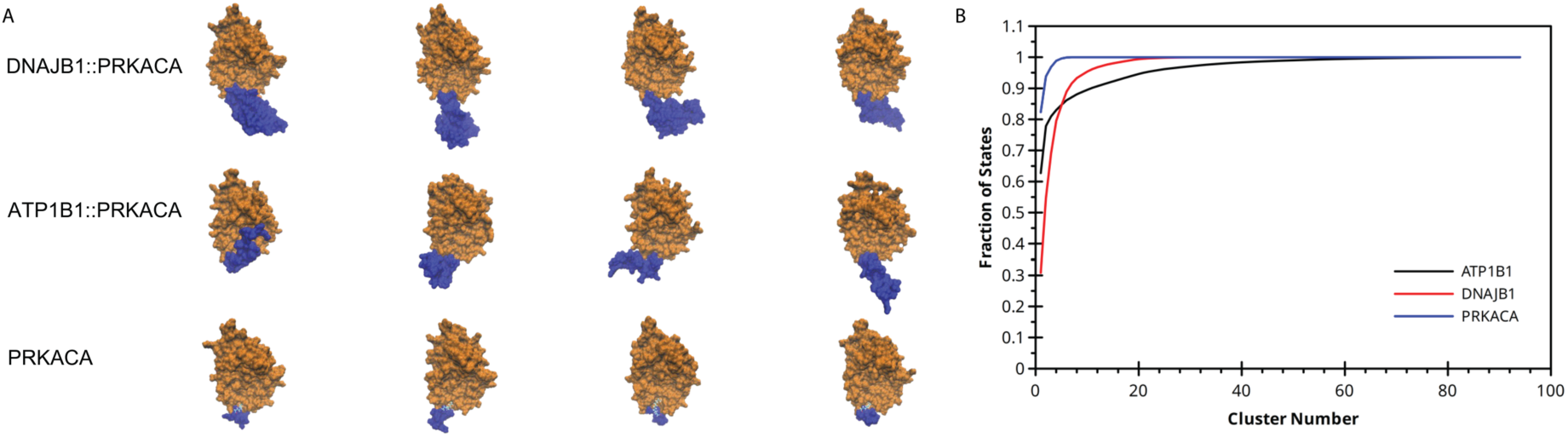
A) Space-filling models of the most frequent conformations of each kinase. Exon 1 encoded N-terminal domains are in blue, while the remainder of the structures are in orange. B) Cluster analysis of the entire conformational space for N-terminal domain of each kinase. DNAJB1::PRKACA and ATP1B1::PRKACA sample many more conformational states than PRKACA.

**Supplemental Figure 5.**
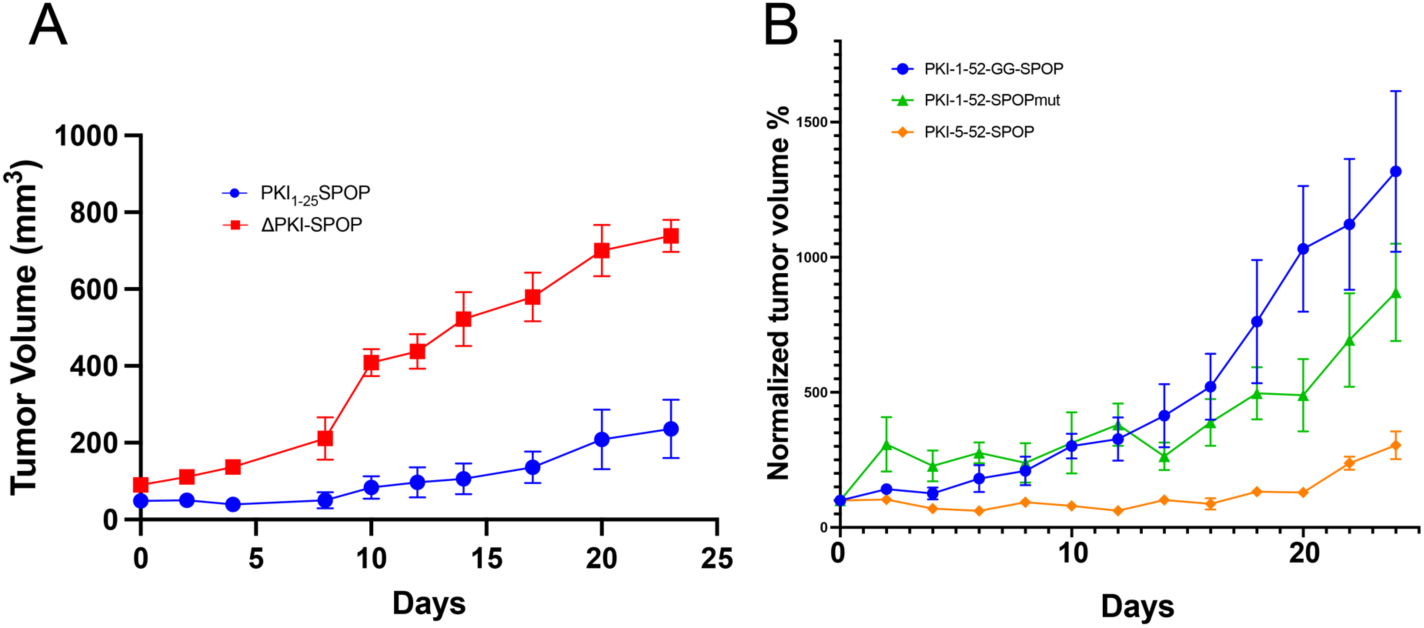
(Figure legend needed): Effect of variants of PKI-SPOP on growth of FLC tumors in mice. A) Quantification of the effects of PKI_1-25_SPOP and ΔPKI-SPOP on FL tumor volume. B) Quantification of the effects of PKI_5-52_SPOP, PKI(_1-52_)SPOPmutant and PKI(GG)-SPOP on normalized FLC tumor volume (n= 5 mice per condition).

**Supplemental figure 6:**
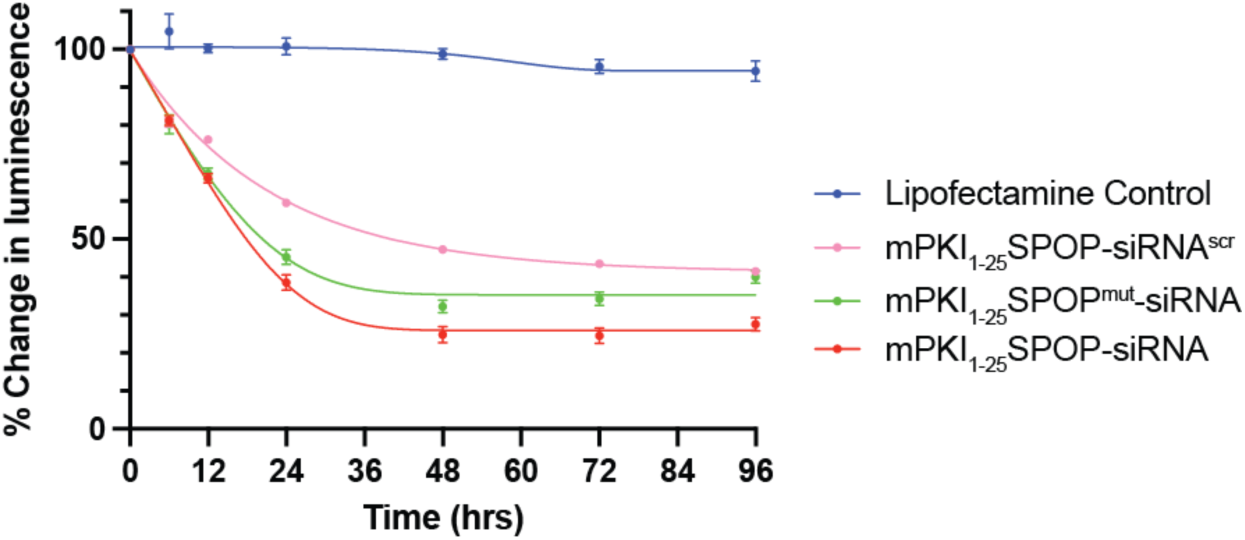
Time course of degrader and antisense. Huh7.5 cells were transduced to stably express DNAJB1::PRKACA-nanoluciferase from the EIF1α promoter. mRNA encoding the biomodal degrader/siRNA (PKI_1-25_SPOP-siRNA, red), or PKI_1-25_SPOP^mut^-siRNA (which inactivated SPOP green), or PKI_1-25_SPOP, PKI_1-25_SPOP-siRNA^scr^ (with the siRNA sequence scrambled), and PKI_1-25_SPOP^mut^-siRNA^scr^ was transcribed in vitro, incubated with the RNA/DNA hybrid and then transfected with lipofectamine into the Huh7.5 cells.

